# Spatio-temporal Coordination of Active Deformation Forces and Wnt / Hippo-Yap Signaling in *Hydra* Regeneration

**DOI:** 10.1101/2023.09.18.558226

**Authors:** Ryo Suzuki, Tetsuya Hiraiwa, Anja Tursch, Stefanie Höger, Kentaro Hayashi, Suat Özbek, Thomas W. Holstein, Motomu Tanaka

## Abstract

Ample evidence suggests that Wnt signaling and tissue deformation are key determinants for pattern formation in animals. The coordination of these biochemical and biomechanical spatio-temporal asymmetries is often unknown or controversial. We investigated this relationship by studying regeneration in the freshwater polyp *Hydra*. In both reaggregates of dissociated cells and tissue regenerates, we found significant tissue contraction waves and upregulation of Wnt signaling. Applying a simple mechanical model to the mode analysis of the active deformations, we quantitatively defined the phase reversal of size change and axial deformation in those oscillations as the time point of “biomechanical” symmetry breaking. Moreover, overexpression and inhibition of canonical Wnt signaling modulated the timing of this biomechanical symmetry breaking. A direct comparison with the RNAseq data indicates that the biomechanical symmetry breaking occurs only after the upregulation of canonical Wnt signaling. Further data suggest that biochemical signaling and biomechanical active deformation synergistically stabilize the body axis and hence the following head structure formation by Hippo-Yap signaling. The symmetry breaking mechanism identified here in *Hydra* most likely represents a patterning module that is evolutionary conserved from early metazoan to bilaterian animals.

## Introduction

During development, the *bauplan* of multicellular organisms is initiated by the emergence of asymmetry, which is referred to as symmetry breaking. From the viewpoint of geometry, this is nothing but an irreversible establishment of the body axes. For example, cell-cell interactions define the body axes in *Drosophila* already during oogenesis, while the sperm entry point determines the location of the anterior-posterior and the dorso-ventral axes in *Xenopus* embryos. However, little is known how the intrinsic or extrinsic cues are coordinated in space and time to initiate symmetry breaking and hence axis formation.

The freshwater polyp *Hydra* is widely known for its unlimited capability of regeneration and hence is considered as potentially immortal. These simple animals are radial symmetric and have a gastrula-like *bauplan* with one mouth opening at one end of the body axis (i.e., the hypostome or so-called “head”) and with an adhesive region (i.e., the peduncle with the “foot”) at the opposite end of the body axis. The formation of new heads and feet occurs by budding; *Hydra’s* way of asexual reproduction, or by regeneration, namely after head and/or foot removal or by *de novo* formation from dissociated single cells in reaggregates (1-7). During *de novo* pattern formation, head- and foot-organizing centers are stochastically formed when cells with higher levels of head competence begin to inhibit the formation of new centers in the neighborhood (lateral inhibition). The cells recruit and instruct surrounding cells to participate in formation of the head and body axis. We showed previously that such clusters of 5 – 15 cells (*Φ* ≈ 45 μm) express *Bra1* and *Wnt3* genes and can act as a head-organizing center (8, 9). We also demonstrated that the domains locally expressing *Wnt3* and *Tcf* define head-organizing centers during budding and tissue regeneration, indicating that Wnt signaling plays a central role in axis formation in *Hydra* (9). The emergence of the small domains that are expressing genes and proteins in a position-specific manner can be considered as a “biochemical symmetry breaking”.

The advantage of *Hydra* among various animal regeneration models is that it can undergo regeneration from both a cut piece of tissue (regenerate) and an aggregate of dissociated cells, in which the level of head-organizing competency can be reset to the random state (7-11). Following the extensive studies demonstrating the vital role of the biochemical symmetry break and hence the *de novo* formation of head-organizing centers (8, 9, 12-14), an increasing number of studies suggested that the active deformation of reaggregates of dissociated cells (Fig. **1a** left) (8-11, 15) and regenerates (Fig. **1a** right) (16, 17) plays a key role in *Hydra* regeneration. Ott and co-workers monitored the deformation of regenerates embedded in agar gels (16) or in droplets (18), and proposed that the morphological symmetry breaking occurs at the time point when the frequency of periodic deformation becomes higher. Keren and co-workers monitored the orientation of actin fibers in regenerates taken from different body parts, and suggested that the actin fiber orientation inherited from the parent animal determines the body axis after regeneration (19). Collins and co-workers have shed light on the mechanical aspects of cell sorting (10) and mouth opening (20). It was also reported that changes in sucrose concentration in the medium affect the frequency of osmotically driven inflation-burst cycles and *Wnt* expression (21). In the anthozoan polyp *Nematostella vectensis* muscular-hydraulic oscillations were shown to drive larva-polyp morphogenesis (22). While these data suggest a correlation of biochemical and biomechanical cues during regeneration, the coordination and transduction of biochemical and biomechanical cues in space and time remain unclear. In vertebrates, a combination of theoretical modeling and experiments has been used to investigate how the mechanical properties of multicellular assemblies are correlated to generate morphological asymmetry in vitro (23-25). However, a comprehensive biomechanical model of symmetry breaking on the organismal level is still missing.

**Fig. 1:**
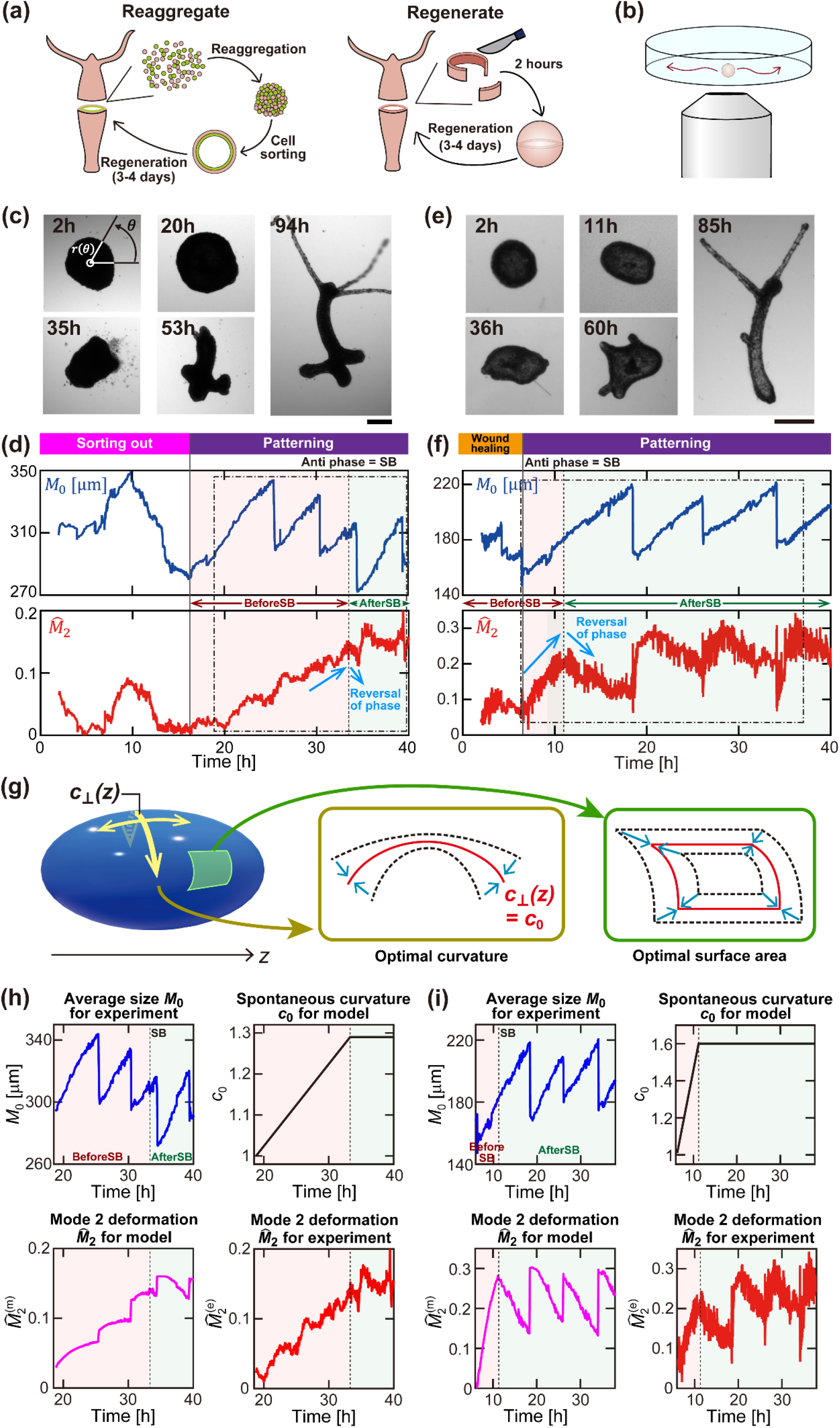
Biomechanical symmetry breaking in *Hydra* reaggregates and regenerates. (a) Preparation of a *Hydra* reaggregate made out of dissociated single cells (left) and a regenerate that is a cut piece of tissue (right). (b) Constraint-free observation of active deformation and motion of reaggregates and regenerates. (c) Snapshot images of a reaggregate over time. Radial distance from the center of mass *r*(*θ, t*) was used for the mode analysis. Scale bar: 300μm. (d) Mode 0 (*M*_0_, blue) and normalized mode 2 (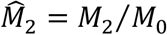, red) of a reaggregate recorded over 40 h. The initial cell sorting out phase (*t* = 0 – 16 h, white background) is followed by the patterning phase. The latter is characterized by the periodic burst-inflation cycles. At the beginning (red background), 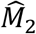 follows the increase in *M*_0_. The time point at which *M*_0_ and 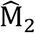 become out of phase with one another (*t* ≈ 33 h; dashed line) is defined as the biomechanical symmetry breaking (SB). After this point (green background), 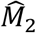 remained higher than 0.15, implying the broken symmetry is sustained. (e) Snapshot images of a regenerate over time. Scale bar: 300μm. (f) *M*_0_ (blue) and 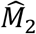 (red) of a regenerate recorded over 40 h. As a regenerate inherits the ecto- and endo-dermal layers from the parental *Hydra*, a wound healing (*t* = 0 – 6 h, white background) is followed by the patterning phase (red/green backgrounds). The biomechanical symmetry breaking is observed at *t* ≈ 11 h (dashed line). (g) Schematic diagram of the theoretical model used to describe biomechanical symmetry breaking. Both reaggregates and regenerates gain the optimal shape by minimizing the total free energy determined by curvature and surface area (red). (h) Model validation using reaggregate data in the patterning phase. Top-left: Experimental *M*_0_ data (*t* = 19 – 40 h) extracted from the dash-dot box in panel (d). Top-right: The model assumes that the spontaneous curvature *c*_0_ increases linearly until SB and remains constant at a fixed value after SB. Bottom-left: Monotonic increase in simulated 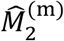 is reversed at SB, and 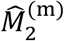 became anti-phase with respect to *M*_0_ . Bottom-right: Direct comparison with the corresponding experimental 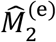. (i) Model validation using regenerate data. Top-left: Experimental *M* data (*t* = 6 – 37 h) extracted from the dash-dot box in panel (f). Otherwise, the calculations and data treatment follow those used for panel (h). See text for further details.

In this study, we monitored the dynamic deformation of *Hydra* reaggregates and regenerates over the entire period of regeneration (Fig. **1**). We ensured that reaggregates and regenerates can freely undergo active deformation and translational/rotational motions with no constraints. We defined biomechanical symmetry breaking by the combination of Fourier mode analysis of active deformation and a simple mechanical model that considers anisotropic spontaneous curvature formation. We also performed a large scale RNAseq analysis of β-catenin and Wnt3 overexpressing animals and compared it with our proteome and transcriptome of *Hydra* head regenerates (26). The direct comparison of deformation patterns and expression dynamics unraveled the coordination of biochemical and biomechanical symmetry breakings. By using transgenic animals and extrinsic factors, we examined how the perturbation on Wnt signaling modulates the timing of biomechanical symmetry breaking and the downstream events like the head structure formation. We also analyzed Hippo-Yap signaling and found that it is involved in the robust orchestration of biochemical and biomechanical signals in regeneration. Our data explain the long-standing question of how biomechanical forces and patterning are linked in *Hydra*, and indicate that the Wnt-Yap interaction found in vertebrates and insects has its roots in the first metazoan animals.

## Results

### Biomechanical symmetry breaking during *Hydra* regeneration

We analyzed *Hydra* regeneration starting from two different initial stages; reaggregates and regenerates (Fig. **1a**). In reaggregates, dissociated cells of *Hydra* gastric tissues first sort out and form ecto- and endoderm before pattern formation starts. In regenerates, gastric tissue with an intact body wall was excised which then regenerates an intact polyp.

Specimens were placed on the glass bottom of a Petri dish with no constraints, which enabled to monitor not only active deformation but also translational and rotational motions during the regeneration process (Fig. **1b**). This is clearly different from previous studies, where specimens were fixed in agar gels or confined in hanging liquid droplets (18), which affects the whole deformation dynamics of the system. To define the biomechanical symmetry breaking geometrically during *Hydra* regeneration, we Fourier transformed the radial distance between the contour and the center of mass *r*(*θ, t*) of the specimens during the whole regeneration process that took about 90 h (Fig. **1c**):

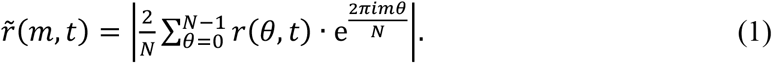

For *m* = 0, the normalization factor is 1/*N*. As *Hydra* tissue pieces exhibit dynamic changes in size and shape during the establishment of the body axis, we analyzed the 0^th^ mode 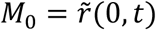 and the normalized 2^nd^ mode 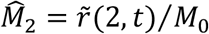 as a function of time. The former reflects the change in averaged radius, while the latter the axis formation.

#### Reaggregate analysis

Figure **1c**shows snapshot images of a *Hydra* reaggregate through the regeneration process. Reaggregates are ultimately a cluster of dissociated cells, which sort out to their origin and form a closed hollow sphere. After *t* = 12 – 18 h, its wall consists of a solid ectoderm and endoderm with an intermediate extracellular matrix (ECM), the mesoglea (1, 8, 10, 11, 27). Figure **1d** shows *M*_0_ and 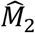 plotted as a function of time. During this “sorting out” phase (white background), neither *M*_0_ nor 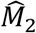 showed any characteristic features. This seems to be in line with the previous studies showing that the ectodermal and endodermal layers form at *t* = 12 – 18 h and the gastric-vascular cavity is becoming functional at *t* = 18 – 24 h (1, 11). After sorting and formation of the two germ layers, we observed saw-tooth like changes in *M*_0_, which coincide with repetitive cycles of burst-inflation (Fig. **1d**, upper panel): the inflation corresponds to the osmosis of the outer medium into the inner gastrovascular cavity, while the burst happens once the sphere cannot withstand the inner pressure (15, 28). Shortly before the reaggregate underwent the third burst at *t* ≈ 34 h, we saw that 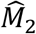 showed a reversal of phase with respect to *M*_0_ at *t* ≈ 33 h indicated by a dotted line in Fig. **1d**. Until this time point, 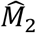 increases monotonically on average and is in phase with the inflation growth of *M*_0_ (Fig. **1d**. Interestingly, this anti-phasic behavior was preserved even during the burst, where rapid increase in 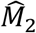was observed, indicating that the reaggregate was axially elongated when it was bursting. After the burst, 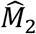exhibited a slow relaxation but the level did not return to its pre-burst value 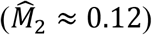 and remained higher than 0.15. The maintenance of the elliptic shape even during the inflation indicates that a stable primordial axis is established when 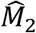 demonstrated a switch from in-phase (red background) to anti-phase (green background) behavior with respect to *M*_0_. Hence, we define the switch to anti-phase behavior at *t* ≈ 33 h as the time point of biomechanical symmetry breaking in reaggregates which is about 15 – 21 h after germ layer restoration.

#### Regenerate analysis

As shown in Fig. **1e**, a piece of tissue with intact ectoderm and endoderm was cut out of the gastric region that regenerated an intact *Hydra* polyp in *t* ≈ 60 – 80 h. Compared to reaggregates from dissociated cells, the regeneration from cut pieces of tissue is about 18 – 24 h faster because regenerates already inherit an intact body wall with ectoderm, endoderm, and mesoglea. By comparison, in reaggregates, dissociated cells have not only lost their initial polarity but also must form both germ layers by a cell sorting and synthesis of a new ECM (mesoglea). The absence of cell sorting in regenerates enabled us to focus sharply on the regeneration dynamics from injury to patterning (29, 30). Depending on the axial position, wound healing (Fig. **1f**, white background) is followed by the patterning phase starting earliest after *t* ≈ 6 h. Indeed, the characteristic dynamic deformations of a regenerate were very similar to that of a reaggregate during the patterning phase. Figure **1f** (upper panel) shows the repetitive cycles of burst-inflation from the very beginning. A distinct switching from in-phase (red background) to anti-phase (green background) between *M*_0_ and 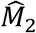 was observed at *t* ≈ 11 h (Fig. **1f**, black dashed line), followed by an anti-phasic increase in 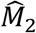 at the time point of burst (*t* ≈ 19 h) and a slow relaxation and maintenance above 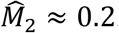. These data coincide with features from the reaggregate experiment, where the symmetry breaking of *Hydra* reaggregates was observed at *t* ≈ 33 h (Fig. **1d**). The difference in the timing of symmetry breaking Δ*t* ≈ 24 h agrees well with the time necessary to create a hollow structure surrounded by ectodermal and endodermal layers in the reaggregate condition and is similar to the duration of the cell sorting phase, *t* = 18 – 24 h (Fig. **1d**). This finding implies that the biomechanical symmetry breaking in *Hydra* regenerates takes about 9 – 12 h after the onset of regeneration including wound healing.

### Theoretical model of biomechanical symmetry breaking

How can the axial deformation 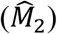 start to decrease in the middle of a size inflating process (*M*_0_)? In addition, why should the abrupt decrease in size (*M*_0_) be accompanied by an anti-phasic increase in the axial deformation 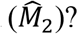? To understand this biomechanical symmetry breaking phenomenon caused by active deformation, we theoretically modeled how an elastic ellipsoid gains its optimal spontaneous curvature and surface area (Fig. **1g**). The assumption of this model is that an anisotropic contraction acts only in the direction of the principal curvature and perpendicular to the major axis of the ellipsoid *z, C*_⊥_(*z*). Under conditions of a fixed volume, the system minimizes energy cost that is caused by deviation from the spontaneous curvature *c*_0_ and optimal surface area, which is given by the potential 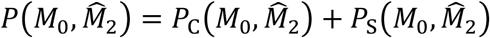, where *P*_C_ and *P*_S_ are the contributions of curvature elasticity and area elasticity of the ellipsoid surface, respectively (see Materials and Methods, Supporting Information **S1** for details). To validate our model, we took the experimental *M*_0_ data from reaggregates (*t* = 19 – 40 h, Fig. **1h**, top left) and regenerates (*t* = 6 – 37 h, Fig. **1i**, top left), and examined if our model could reproduce the experimental 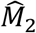 values. We assumed that the volume of a *Hydra* is fixed at each time point, and it is determined to be consistent with the observed *M*_0_. For the sake of simplicity, we implemented in the model that the maturation of spontaneous curvature *c*_0_ increases linearly before the symmetry breaking (Figs. **1h**, and **i**, top right). At the point of symmetry breaking, *c*_0_ reaches a certain value and remains constant afterwards corresponding to the establishment of a fixed axis. As shown in Figs. **1h** and **1i**, 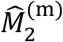 calculated by minimization of 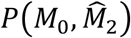 at each time point recapitulated the experimentally determined 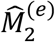 .

It should be noted that our model calculation is quantitative because the previously reported mechanical parameters of *Hydra*, such as coefficient for bending and area rigidity (17, 31), were used in the simulations (see Materials and Methods, Supporting Information **S2** for details). Altogether the current data suggest that the emergence of symmetry breaking requires active deformation, which over time perturbs and disrupts the initial symmetry in the system (32).

### Roles of active force generation by the actomyosin complex

The main component that contributes to active deformation in a *Hydra* is the actomyosin complex. Figure **2a** shows the fluorescence microscopy image of an adult *Hydra* (left) and its zoom up (middle), stained with phalloidin 488. The orthogonal grid-like features originate from the overlay of actin filaments in the ectoderm and endoderm. Actin filaments in ectoderm are aligned parallel to the oral-aboral body axis, while those in endoderm are perpendicular. Figure **2b** represents the fluorescence image of a tissue regenerate taken at *t* = 2 h and an analysis of the orientation of phalloidin 488 stained actin filaments. The orientation order of actin filaments was quantitatively assessed by the orthogonal order parameter Φ = cos 4*φ*_ij_,where *φ*_ij_ is the angle between the i-th and j-th filament structure in regenerates and reaggregates (Fig. **2c**). Here, the ensemble average ⟨Φ⟩ = 1 denotes filaments crossing perpendicularly to make a perfect mesh structure, whereas ⟨Φ⟩ = 0 means that the filaments are completely random.

**Fig. 2:**
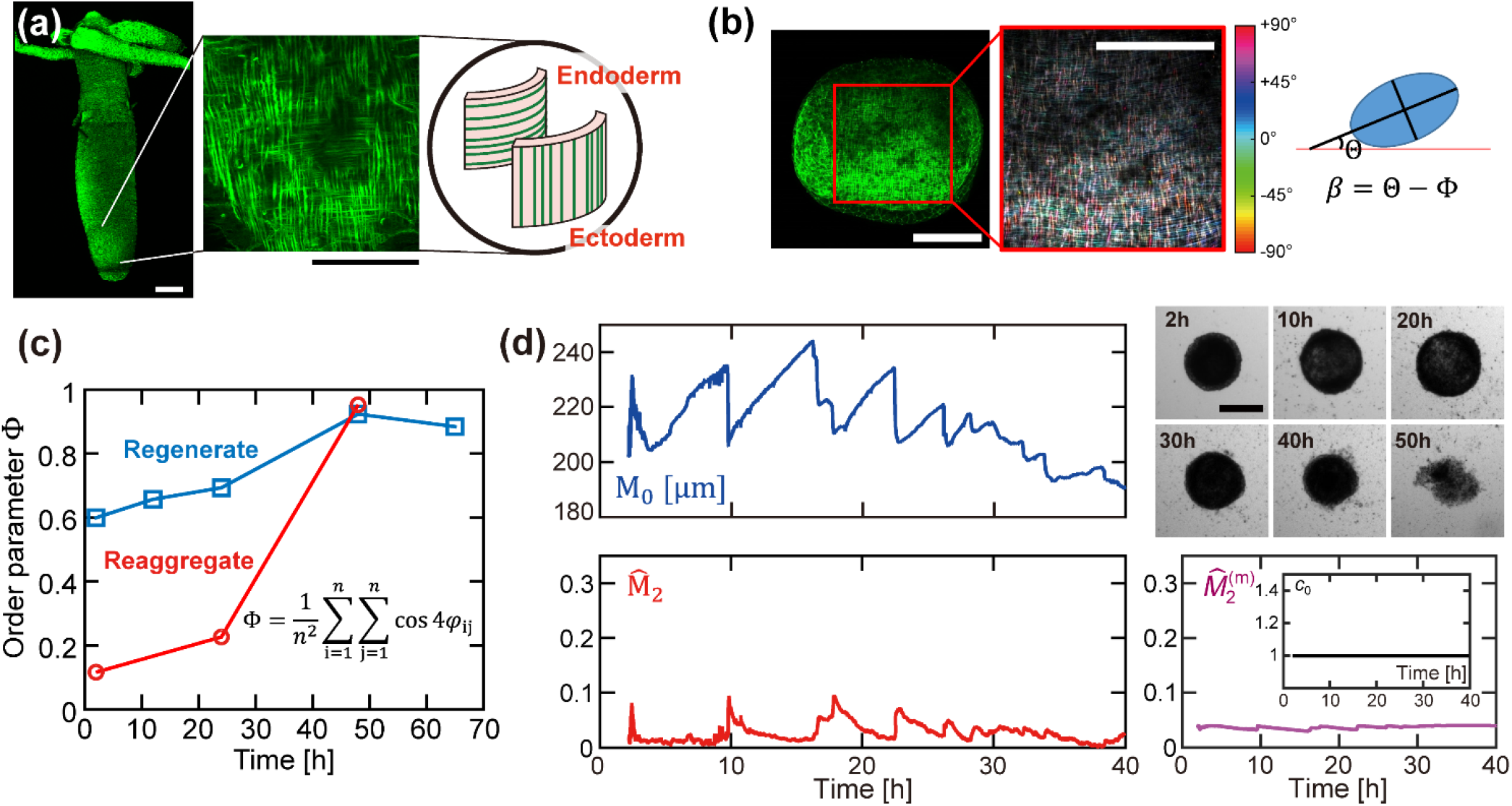
Ordering of actin cytoskeletons during *Hydra* regeneration of a tissue regenerate. (a) Actin cytoskeletons of an adult *Hydra* stained with phalloidin 488. A zoom-up image exhibits mesh-like actin structure. Note that actin fibers in ectoderm are aligned parallel, while those in endoderm are aligned perpendicular to the oral-aboral body axis. (b) Distribution of actin orientation in the region of interest. The color indicates the angle of individual actin fibers. (c) Orthogonal order parameter of actin fibers in a rectangular grid, Φ = cos 4*φ*_ij_. Growing actin fibers in a reaggregate (red) gains orientation order over time, while those in a regenerate sustained an ordered actin structure inherited from the parental animal. (d) *M*_0_ (blue) and 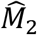 (red) of a regenerate treated with 20 μM myosin II inhibitor blebbistatin, recorded over 40 h (left). The regenerate is still capable of the repeated burst-inflation cycles by osmosis, but 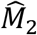 remains close to 0 at all times and no anti-phasic behavior is observed. As seen in the snapshot images (top-right), the regenerate disintegrated. Recapturing the lack of deformation 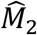 for the blebbistatin treated regenerate with the mechanical model (bottom-right), using a constant *c*_0_ = 1 (inset). Scale bars: (a,d) 300μm, (b) 100μm.

Our analysis is in good agreement with a previous report, showing that the architecture of the actin-network in epithelial cells accompanies body axis formation (19). We found, however, different dynamics in the formation of the final mesh structure in reaggregates and tissue regenerates. This can be seen from the average order parameters calculated for the reaggregate and the regenerate, ⟨Φ_reag(24h)_⟩ = 0.23 and ⟨Φ_rege(24h)_⟩ = 0.70, respectively. This difference can be explained by the fact that a reaggregate consists at *t* = 0 h of a blend of ectodermal and endodermal cells with no ordered cytoskeletons (11). Over time, patches of mesh-like actin filaments gradually grew (Fig. **S1**), reaching *φ*_*reag*_ ≈ 1 at *t* ≈ 50 h (Fig. **2c**), which is similar to regenerates, that already start at a high order parameter value. Hence, in the *de novo* system with no inherited actin order, the maturation of the orthogonal actomyosin complexes does not guide the biomechanical symmetry breaking, as argued by Ref. (19).

To unravel the interplay between actomyosin maturation and biomechanical symmetry breaking, we treated a regenerate after the formation of closed sphere with blebbistatin, which selectively inhibits non-muscle myosin II ATPase activity (33). We used a lower dose (20 μM) compared to Ref. (19) and analyzed how the dynamic deformation was modulated. The regenerate did not lose the capability to undergo the osmosis driven, repeated cycles of burst-inflation (Fig. **2d**, upper panel). Although the burst was accompanied by a rapid increase in 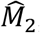, it did not stay at a high level (≥ 0.15) but went down almost to the baseline (Fig. **2d**, lower panel) implying that the regenerate became spherical, because no contractile force can be generated due to the myosin II inhibition. Consequently, the blebbistatin-treated regenerate was not able to establish a stable primordial axis and hence could not undergo regeneration. After 40 – 50 h, we observed the disintegration of the regenerate. The lack of anisotropic tissue deformation in our blebbistatin experiments can be translated within the framework of the mechanical model as a maturation failure of the spontaneous curvature *c*_0._ Using the *M*_0_ data from Fig. **2d** and keeping *c*_0_ close to 1, the experimental results, no anti-phasic behavior between the two modes and low 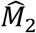 values, were quantitatively reproduced (Fig. **2d**). The obtained results demonstrated that the maturation of ordered actomyosin complexes and thus spontaneous curvature *c*_0_ is necessary for the generation of active deformation force driving biomechanical symmetry breaking.

### Wnt signaling acts upstream of biomechanical symmetry breaking

We next examined whether the biomechanical symmetry breaking occurs before or after the “biochemical symmetry breaking”. Previous work revealed a strong upregulation of Wnt3 and β-catenin within the first three hours after injury of regenerating head (8, 34) and foot tissue (29, 30, 35). This early expression of *Wnts* is part of a generic wound response, which becomes head-specific only *t* ≈ 6 – 12 h after injury and leads to a proper regenerating head organizer while Wnt signaling is downregulated in the presumptive foot (29, 30, 35, 36). Figure **3a** shows an ISH of Wnt3 in regenerates that originated from a piece of tissue that was cut out of the gastric region (Fig. **1e**). Such pieces exhibit an oral, aboral, and lateral wound side which rapidly close, form a hollow sphere within *t* ≈ 2 h, and regenerate to intact polyps in *t* ≈ 60 – 80 h (Figs. **1e, 3a**). During sphere formation, the injured tissue exhibits a strong *Wnt* expression. From *t* ≈ 8 h on, *Wnt* expression diminishes between the two poles of the sphere. The *Wnt* pattern in such regenerates is very similar to excised rings of gastric tissue forming a head at one end and the foot at the other end (29). At *t* ≈ 14 -26 h, *Wnt* expression is sustained only at one pole, from which the future head emerges, while the opposite pole forms the future foot. These ISH data clearly indicate that generic Wnt signaling precedes biomechanical symmetry breaking in the regeneration process. However, they also demonstrate that a stable and head specific *Wnt* expression pattern only arises at *t* ≈ 8 -14 h, which surprisingly correlates with the time of biomechanical symmetry breaking in our regenerates (Fig. **1f**).

**Fig. 3:**
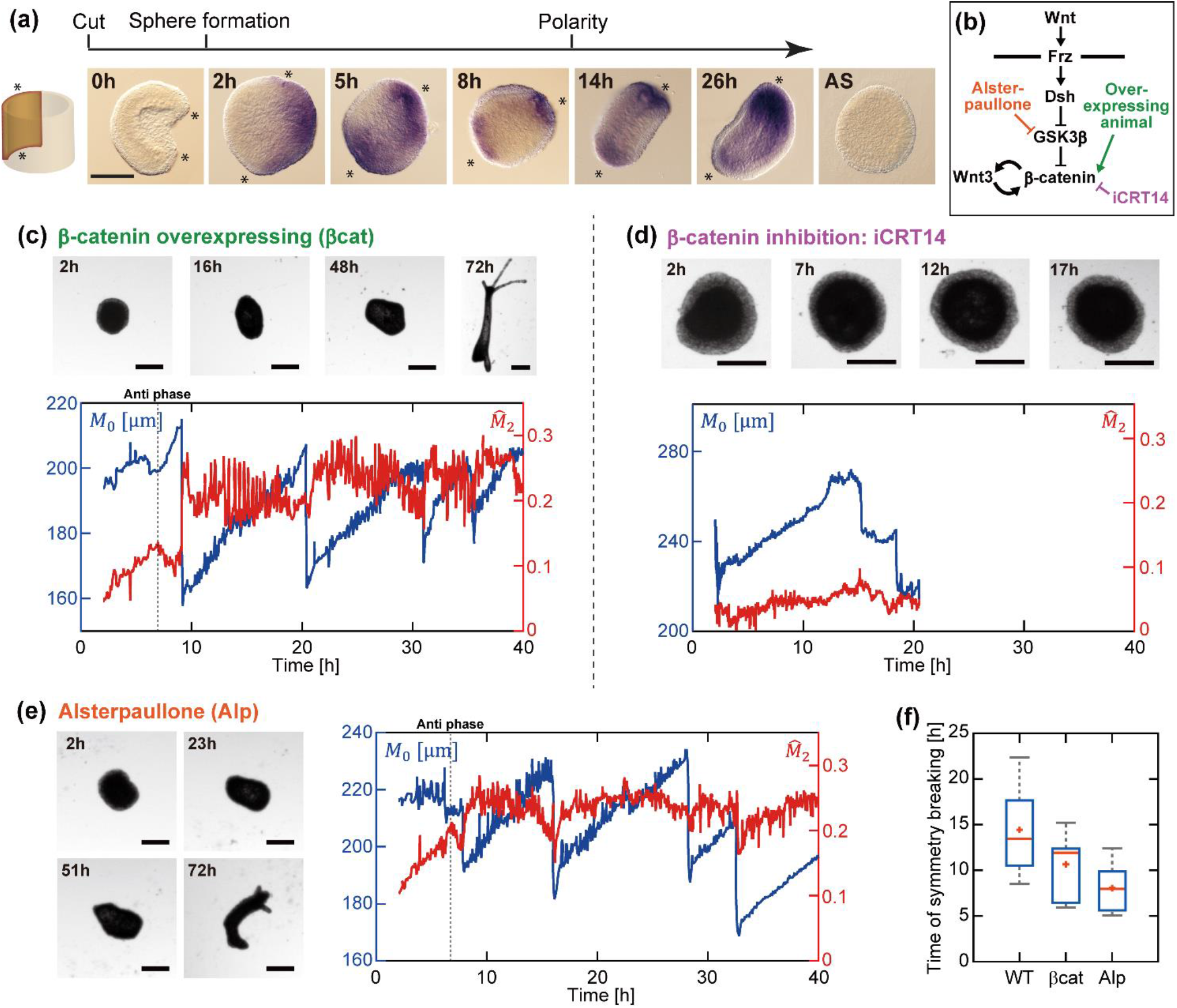
Perturbation on Wnt signaling modulates biomechanical symmetry breaking. (a) ISH of small pieces indicates an early upregulation at the wound side and the presumptive head and foot tissue (asterisks), which becomes focused to presumptive head tissue indicated by polarized Wnt expression between *t =* 24 – 36 h after cutting. (b) Wnt signaling pathway. (c) Snapshot images of a regenerate from a β-catenin overexpressing animal. *M*_0_ (blue) and 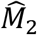 (red) of a regenerate overexpressing β-catenin, exhibiting a biomechanical symmetry breaking at *t* ≈ 7 h. (d) Snapshot images of a regenerate treated with β-catenin inhibitor, iCRT14 (20 μM). After 20h, the regenerate disintegrates and does not regenerate. *M*_0_ (blue) and 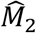 (red) of a regenerate treated with iCRT14 recorded over 40 h. (e) Snapshot images of an Alsterpaullone (Alp) -treated regenerate. A regenerate treated with Alp underwent a biomechanical symmetry breaking at t ≈ 6 h. (f) Comparison of the time of biomechanical symmetry breaking for wild-type (n = 10), β-catenin overexpressing animal (n = 6) and Alsterpaullone treated (n = 6) regenerates. Scale bars: (a) 200μm, (c-e) 300μm.

The correlation of the kinetics of biomechanical symmetry breaking (Fig. **1a**) with “biochemical symmetry breaking” (Fig. **3a**) suggests that mechanical cues and the formation of a stable Wnt-related head organizer are linked. To verify if the perturbation of Wnt signaling has an impact on the biomechanical dynamics described above, we next used transgenic and pharmacological approaches to perturb Wnt signaling (Fig. **3b**): (i) transgenic animals overexpressing β-catenin (14), (ii) iCRT14 treated animals in which β-catenin was inhibited (36, 37), and (iii) Alsterpaullone (Alp) treated animals with global β-catenin activation by GSK3 inhibition (13, 29, 38, 39). Figure **3c** represents the snapshot images of the regenerate produced from a β-catenin overexpressing animal. The regenerate underwent a biomechanical symmetry breaking at an earlier time point (*t* ≈ 7 h) compared to a wild type regenerate (*t* ≈ 11 h). This is reasonable, as β-catenin overexpressing animals possess a high level of head activation, often generating multiple heads and feet (40). Conversely, the wild type regenerate treated with the β-catenin inhibitor iCRT14 (20 μM) (36, 37), resulted in no regeneration (Fig. **3d**). In fact, the characteristic burst-inflation cycles driven by contraction of actomyosin complexes during regeneration were not observed. Since β-catenin does not only act on Wnt signaling but also controls cell adhesion via the actin cytoskeletons (41), we extrinsically manipulated the Wnt signaling by using Alp as GSK3-β inhibitor (13, 36, 42). As shown in Fig. **3e**, the indirect activation of Wnt by the treatment of 0.05 μM Alp on the wild type regenerate resulted in the emergence of biomechanical symmetry breaking at a distinctly earlier time point (*t* ≈ 6 h), similar to β-catenin overexpressing animals. These data imply that perturbations of Wnt signaling can significantly modulate biomechanical symmetry breaking as much as half the time necessary for a wild type regenerate (Fig. **3f**).

### Perturbations of Wnt signaling modulate head structure formation

The interplay of mechanical and biochemical signals finally modulates various downstream events such as head structure formation and tentacle formation. We therefore analyzed how perturbations of β-catenin and Wnt signaling affect the formation of *Hydra’s* mouth, the ultimate terminus of the aboral-oral body axis. Fig. **4a** shows the time progression of *M*_0_ of a wild type animal over the whole regeneration process. In the snapshot images taken at the burst events, the positions of burst (red), future head structure (green), and future foot (yellow) are indicated by symbols. Remarkably, the position of burst and that of the future head match only at *t* ≈ 60 h, when the frequency and amplitude of burst-inflation cycle became smaller.

**Fig. 4:**
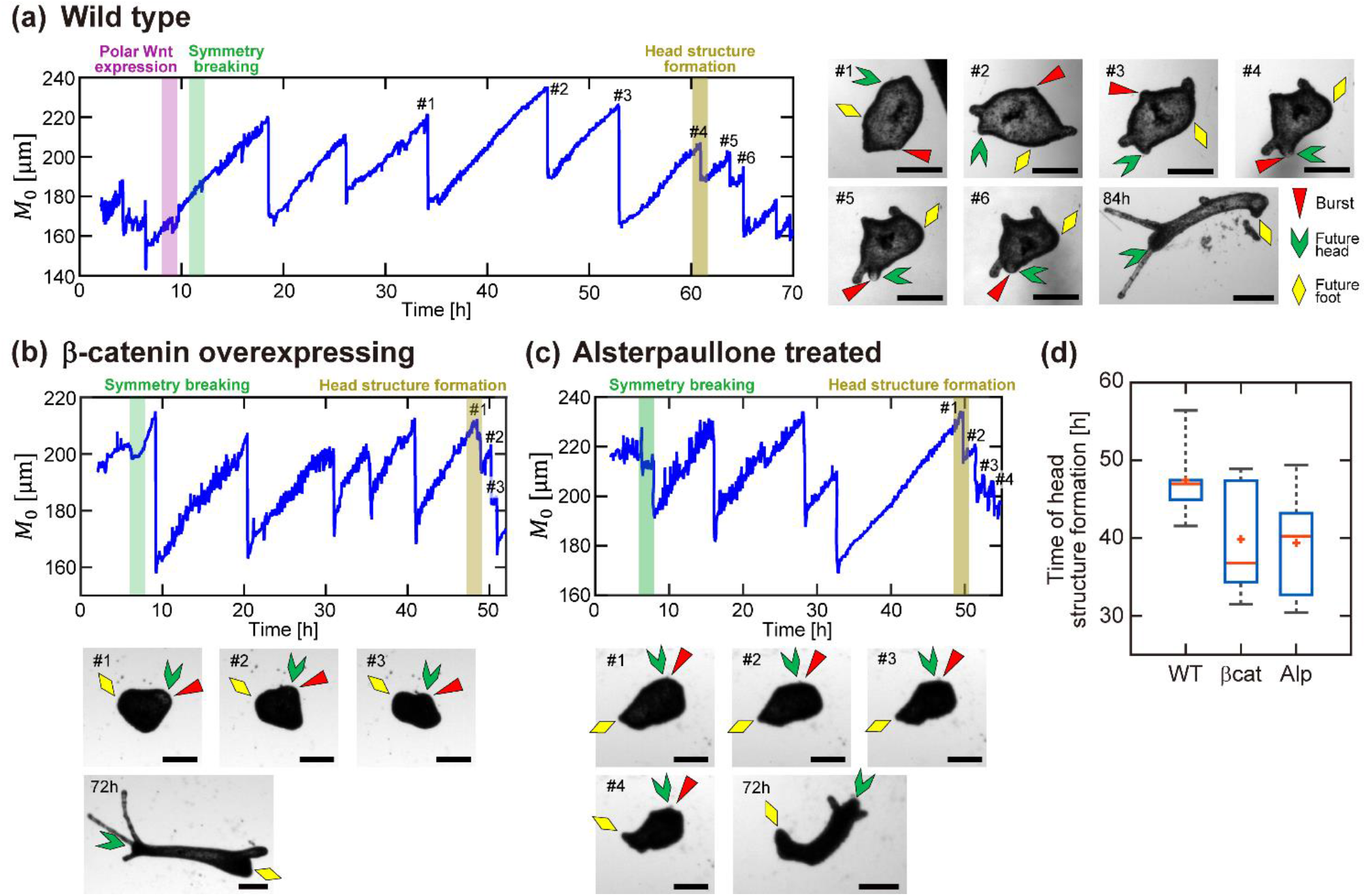
Perturbation on Wnt signaling modulates downstream events (head structure formation). Correspondence of the positions of bursts (red), future head structures (green) and future feet (yellow) monitored for (a) regenerate from a wild type *Hydra*, (b) regenerate from a β-catenin overexpressing animal and (c) regenerate treated with Alp. The number given to each image coincides with the number indicated in each *M*_0_ vs *t* plot. The colocalization of the burst point and the future head was observed about (a) 61 h, (b) 49 h, and (c) 49 h after the biomechanical symmetry breaking. (d) Comparison of the time of head structure formation for wild-type (n = 10), β-catenin overexpressing animal (n = 6) and Alsterpaullone treated (n = 6) regenerates. All scale bars are 300 μm.

This time point, at which the amplitude and frequency of the size exhibit a significant change, was defined as the point of symmetry breaking in previous studies (16, 17, 43, 44). However, our data suggest that this definition should be revised, because the final head formation and head morphogenesis is a downstream event after the establishment of the body axis by “biochemical” and “biomechanical” symmetry breaking (see Discussion). More recently, fluorescently labeled microbeads have been injected into regenerates embedded in agarose gels to observe the positions of osmotic bursts over time (45, 46). It was reported that the change in size oscillation frequency coincides with the formation of head structures, but without clearly defining the time point of symmetry breaking (45, 46). Our definition, based on the theoretical model, enables us to disentangle the head structure formation and the biomechanical symmetry breaking quantitatively. Our data indicate that the formation of head structure takes place *t* ≈ 40 – 45 h after the polar *Wnt* expression and *t* ≈ 35 – 40 h after the biomechanical symmetry breaking. This is consistent with reaggregates where the head structure was formed about 48 h after the onset of *Wnt3* expression (8).

In case of the regenerate prepared from a β-catenin overexpressing animal (Fig. **4b**), the head structure was formed earlier, *t* ≈ 40 h after the biomechanical symmetry breaking. The positions of head and burst were co-localized during the small burst-inflation cycles that follow afterwards. The head structure formation at a much earlier time point compared to the wild type is consistent with the promotion of head-organizing centers in β-catenin overexpressing animal. The same tendency was observed for the wild type regenerate treated with Alp (Fig. **4c**), suggesting that the activation of Wnt signaling leads to the head structure formation at an earlier time point; approx. 40 h after the biomechanical symmetry breaking. Although there are differences in individual regenerates, the general tendency that perturbations of β-catenin (and so of Wnt signaling) led to faster head structure formation holds, resulting in a maximum of nearly half the time necessary for wild type regenerates (Fig. **4d**).

### Hippo-Yap signaling acts during symmetry breaking in *Hydra* head regeneration

Our experiments on the interaction of biochemical and biomechanical signals indicate a temporal precedence of β-catenin and Wnt signals in the regeneration of the *Hydra* head organizer. This does not preclude the release of a cascade of biochemical signals as an “injury signal” at the beginning of regeneration (29, 35). To decipher how the biochemical and biomechanical signals are coordinated in *Hydra* head organizer formation during regeneration, we have tried to identify factors that could link biochemical and biomechanical signals. To this end, we analyzed the transcriptomes of regenerating tip tissue from head regenerates of wildtype animals (26) and strains overexpressing *Wnt3* or *β-catenin* and forming numerous head-related structures (Fig. **5a**). Transcripts that encode signals involved in mechano-transduction should be expressed during regeneration and they should also be up- or downregulated in transgenic animals that are constitutively overexpressing members of the Wnt-pathway. Here, we identified the Yap/Hippo signaling as the major pathway linking biomechanical and biochemical processes (Fig. **5b**). Additionally, we analyzed *β-catenin*- and *Wnt3* overexpressing animals, which exhibits numerous head-related structures (i.e., tentacles), but only *Wnt3* overexpressing animals form well defined ectopic body axes (Fig. **5a**).

**Fig. 5:**
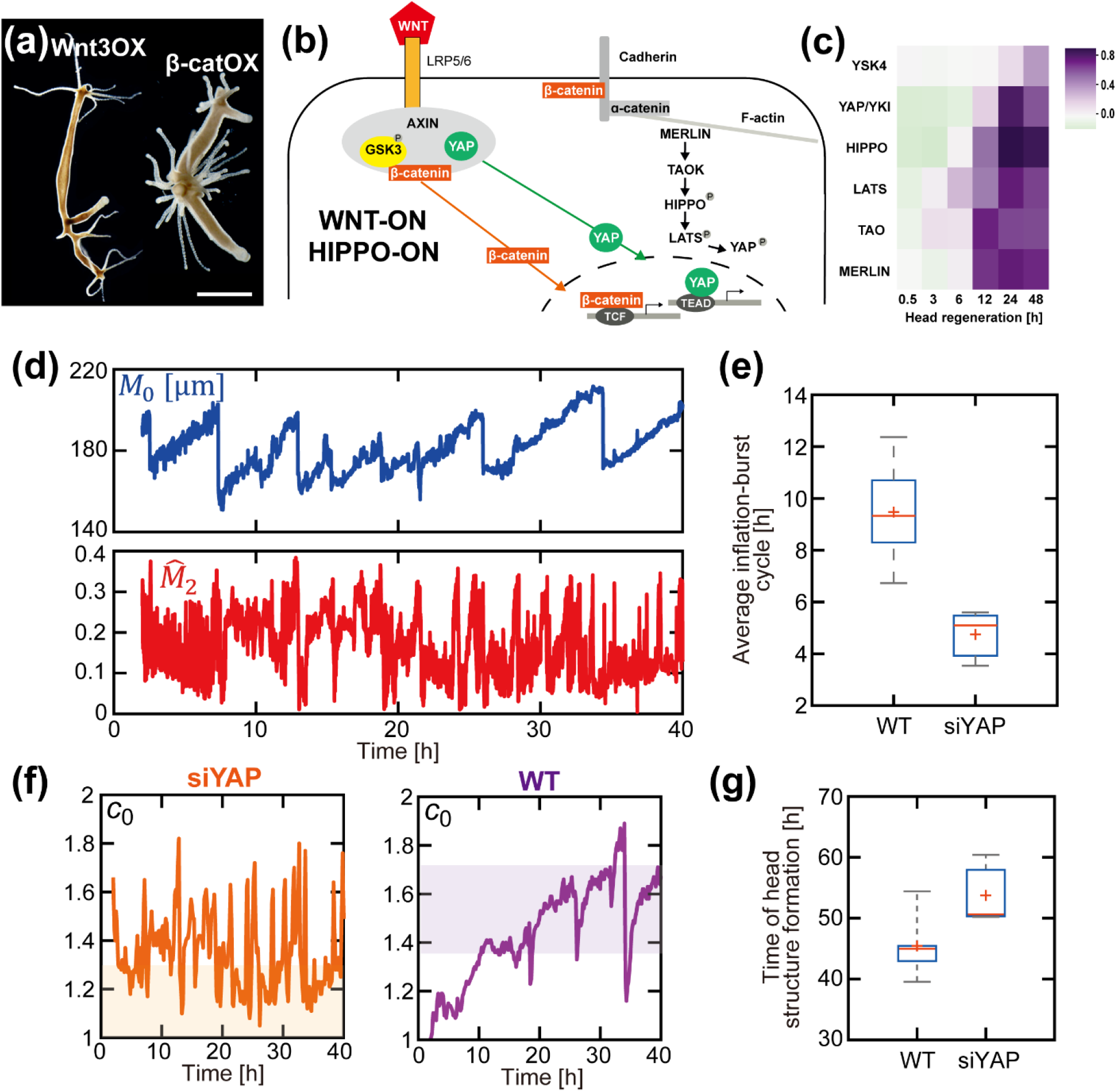
Hippo-Yap signaling in Hydra head regeneration, Wnt3 or β-catenin overexpressing polyps and regenerates. (a) Phenotypes of polyps overexpressing β-catenin (β-catenin::β-catenin-GFP) or Wnt3 (Act::RFP-Act::Wnt3). (b) Scheme showing activated canonical Wnt signaling with nuclear β-catenin and the Hippo pathway with nuclear Yap. (c) Heat map of gene expression levels of selected members of the Hippo-Yap pathway in regenerating tip tissue at the side of head formation (*t* = 0.5 – 48 h); shown by log2-fold changes (blue indicates upregulated and red downregulated genes). (d) *M*_0_ (blue) and 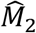 (red) of a siYAP treated regenerate. Inflation-burst cycle becomes faster and active deformation is highly fluctuating. (e) Comparison of inflation-burst cycle between wildtype and siYAP regenerates. (f) Spontaneous curvature *c*_0_ calculated from experimentally obtained *M*_0_ and 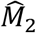, via the model. Unstable and low *c*_0_ values are seen for the siYAP tissue (left), while the wildtype tissue shows a gradual maturation of *c*_0_ and is stable at high *c*_0_ values (right). (g) Time of head structure formation is also affected by Yap.

Based on our proteome/transcriptome analysis (26), we established a list of transcripts that included a total of 15,244 sequences that were analyzed by DESeq2 and hierarchical clustering (Supporting Information **4**). Based on the similarity of their regulatory patterns, we identified 15 clusters (Supporting Fig. **S3**). There was a sharp boundary between clusters during early regeneration (*t* ≈ 0.5 – 12 h) and late regeneration (*t* ≈ 12 – 48 h). Early clusters largely correspond to post injury events before biomechanical symmetry breaking while later clusters were specific for pattern formation and cell differentiation (Supporting Fig. **S3**). Members of the Wnt pathway became activated concomitantly with the onset of regeneration. Some clusters were also different between the *β-catenin* and *Wnt3* overexpressing animals, indicating distinct levels in the regulation of Wnt signaling during regeneration, despite autocatalytic feedback loops (14).

A deeper analysis of regeneration specific genes revealed that the Hippo-Yap pathway was strongly upregulated at *t* ≈ 12 h of head regeneration suggesting that it is a major pathway linking biomechanical and biochemical processes (Fig. **5b, c**). Hippo-Yap signaling acts in size control and mechano-transduction during embryogenesis and cancer (47, 48). The core of the Hippo-Yap pathway consists of a cascade of protein kinases (49, 50) which is evolutionary conserved from early metazoans to insects and mammals (51-53). Activated Hippo finally phosphorylates Yap that is a major factor of mechano-transduction (54-56). Biomechanical input to the Hippo-Yap pathway can occur by Merlin/NF2 (51), which is linked to the actin cytoskeleton and cell adhesion (Fig. **5b**) (57). Our data show that transcripts encoding Tao and Wts/Lats are enriched already *t* ≈ 3 – 6 h after cutting (Fig. **5c**). These transcripts and those encoding for Hippo, Merlin, and Yap were all strongly enriched at *t* ≈ 12 h and up to 48 h after cutting. These data clearly emphasize the importance of Hippo-Yap signaling for *Hydra* regeneration.

Because Yap acts as the mechano-sensing nuclear effector of the Hippo-Yap pathway (54, 58, 59), we analyzed its function in wild type *Hydra* and in our regeneration assay (Fig. **1a**). When we silenced *Yap* in intact animals by the siRNA approach (37, 60), we found related to recent work an increase of bud formation (53).

However, when we tested our regenerates, we surprisingly found that siYAP treated gastric tissue pieces regenerated quite poorly (27%). The analysis of the inflation-burst cycles of such regenerates demonstrates that the inflation-burst frequency was approximately two times shorter for regenerating siYAP tissue compared to that from wildtype tissue (Fig. **5d,e**). The rapid inflation-burst cycles of siYAP-treated regenerates and the low regenerative capacity suggest that the intercellular tension of siYAP-treated regenerates is very low compared to wildtype or β-catenin/Wnt-overexpressing tissue. This is also consistent with siYAP knockdowns showing an increase in the number of buds in the budding region. This tissue has a higher intercellular tension than the upper gastric region (61), so that a loss of Yap could contribute to the softening of the tissue (see Discussion). Moreover, the highly fluctuating 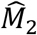 of the siYAP tissue (Fig. **5d**) hinders the specification of the time point of biomechanical symmetry breaking, which suggests that the siYAP tissue is unable to establish a stable primordial body axis in the early time regime of the regeneration process. This can be translated, in the context of the model, as the inability of the siYAP tissue to develop anisotropic deformation and hence maturation of spontaneous curvature *c*_0_. To check this aspect, we used the mechanical model to calculated *c*_0_ from the experimentally obtained *M*_0_ and 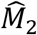. Figure **5f** shows the calculated *c*_0_ for siYAP and wildtype regenerates. For the siYAP case, similar to the behavior of 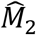, *c*_0_ is unstable and spends the majority of the time at low values close to 1, which indicates that it is absent of anisotropic deformation necessary for biomechanical symmetry breaking. On the contrary, *c*_0_ gradually matures in the wildtype regenerate and stabilizes at a relatively high *c*_0_ value. Consequently, head structure formation is also affected, where the siYAP tissue takes more time for the manifestation of downstream events (Fig. **5g**). Thus, Hippo-Yap signaling assists in the synergistic stabilization of the body axis initiated by Wnt signaling and active deformation.

## DISCUSSION

The regeneration of new heads and feet in *Hydra* is one of the most studied processes in biology and can serve as a paradigm of regeneration biology. The formation of new heads and feet in *Hydra* occurs also in reaggregates from dissociated cells. This process was characterized as *de novo* pattern formation because all positional information is lost during tissue dissociation and the position information for head and foot structures must be restored from a completely stochastic state where all cells initially have the same positional information. The emergence of new signaling centers resulting finally in new heads and feet from an initially random system can be characterized as a “biochemical” symmetry breaking. So far it was considered mainly within the framework of reaction-diffusion theorem (62-64) where an autocatalytic amplification of concentration fluctuations of activators leads to the generation of a pre-pattern for the establishment of a body axis. On the other side, also biomechanical cues have been described to have a major impact on symmetry breaking formation (16, 43, 44), but little is known about the primacy of both mechanisms (19, 21). We have combined both approaches in this study to clearly define the symmetry breaking, both biomechanically and biochemically and how they are orchestrated.

### Re-definition of biomechanical symmetry breaking in *Hydra*

Remarkable roles that biomechanical symmetry breaking play in *Hydra* regeneration have been reported by several studies, but the definition of biomechanical symmetry breaking remains a matter of discussion. Ott and co-workers defined the time point at which the frequency of osmotically-driven inflation-burst cycles of regenerates becomes higher as the symmetry breaking point (16, 17, 43, 44), and the correlation between the inflation-burst cycles and Wnt3 expression was reported by adjusting the osmotic pressure difference between inside and outside of regenerates (21). Keren and co-workers suggested that the axis determination in *Hydra* regenerates is inherited from the parental polyp through the arrangement of the actin cytoskeleton but does not involve symmetry breaking (19). In fact, all the above-mentioned studies have used cut pieces of tissues (regenerates) that inherit not only the ectodermal and endodermal layers but also the ordering of actomyosin complexes from parental *Hydra*. These controversial interpretations indicate that the definition of biomechanical symmetry breaking should be reconsidered from the physical viewpoint.

To decouple the influence of “inherited” actomyosin ordering, we analyzed the active deformation of *Hydra* undergoing regeneration from the *de novo* system (reaggregates) and compared it with that of regenerates in a confinement-free environment. With aid of a mechanical model, we quantitatively defined the biomechanical symmetry breaking as the time point at which modes 0 and 2 starts showing anti-phasic behaviors (Fig. **1**). Such an anti-phasic behavior was common for both reaggregate and regenerate systems. It is notable that the time point of biomechanical symmetry breaking we defined from a mechanical viewpoint is largely different from the time point suggested by previous studies of Ott and co-workers (16, 17, 43, 44). Our time lapse images taken under confinement- and label-free conditions surprisingly demonstrate that the switch of *m* = 0 oscillation frequency they observed coincides with the final formation of the head structure (Fig. **4**), as Wang et al observed by the localization of fluorescent beads (45, 46). This is a downstream event about 40 h after the biomechanical symmetry breaking and overlapping with final mouth formation (Fig. **4a**). Indeed, Ott and co-workers (65) correlated this event with the expression of *ks1*, an epithelial head forming gene whose expression in ectodermal epithelial cells and the evagination of tentacles could be detected at least 2 d after Wnt expression (66). Thus, mouth and final head structures emerge at a very late time point compared to the initial *Wnt* activation (9) and to the biomechanical symmetry breaking point defined here (Fig. **4a**). In terms of pattern formation, the definition of biomechanical symmetry breaking should therefore be revised.

### Biochemical and biomechanical features of symmetry breaking

Previous studies have shown that *Wnt* activation precedes the biomechanical symmetry breaking, which finally results in the final head structure formation. This is evident from the early expression *β-catenin* and *Wnt* genes starting 3 h after head removal or injury (8, 9, 26, 29, 34, 36). Although the early activation of Wnt signaling is part of the generic injury signal stimulated in head and foot regenerates within the first 6 h after injury (29, 30, 35, 36), a wave of head-specific genes becomes upregulated at *t* = 6 – 12 h in presumptive head tissue (Fig. **5c**; Supporting Fig. **S3**). In contrast, biomechanical symmetry breaking emerged at *t* ≈ 11 h (Fig. **1f**). This indicates a primacy of signaling mechanisms. It is also supported by the fact that perturbations of Wnt signaling can significantly modulate the biomechanical symmetry breaking (Figs. **3, 4**).

A critical role in the biomechanical symmetry breaking is played by active tissue deformation generated by the contraction of actomyosin complexes. In fact, the regenerate was not able to establish a stable primordial axis once myosin II was blocked by blebbistatin treatment (Fig. **2**). Previously, Keren and co-workers reported that the body axis in a regenerating *Hydra* tissue piece (regenerate) is “inherited” from parental *Hydra* (19, 67). To disentangle the effect of inherited actomyosin orders, we monitored the dynamic deformation of reaggregates undergoing *de novo* patterning (Fig. **1d**), and found that the timing of biomechanical symmetry breaking in reaggregates was distinctly delayed compared to the regenerates (Fig. **1f**), for two reasons. First, reaggregates sort out to their origin and form a closed hollow sphere (*t* = 0 – 16 h). Second, reaggregates must reestablish orthogonal actomyosin orders to generate active deformation forces. As shown in Fig. **2**, the orthogonal order parameter of actin cytoskeletons is distinctly lower in reaggregates at *t* = 0 – 24 h than in regenerates. This clearly indicates that the organization of actomyosin complexes is necessary for biomechanical symmetry breaking, but not causal.

So far, a combination of mechanistic studies on molecular levels, quantitative measurements, and theoretical modeling that unravel how the spatio-temporal Wnt signaling guides the biomechanical symmetry breaking and regeneration is still missing. A correlation between osmotically driven inflation-burst cycles and *Wnt3* expression was reported suggesting that mechanical forces modulate Wnt signaling (21), but the underlying molecular mechanism remains to be determined. Our data indicate that symmetry breaking during regeneration occurs downstream of Wnt signaling and likely induces changes in further signaling pathways required to pattern the cytoskeleton of *Hydra*. After injury, *Wnt3, Wnt9/10a* and *β-catenin* expression are activated by default as a consequence of generic MAPK phosphorylation to drive tissues into a regeneration competent state (29, 30, 35, 36). Strikingly, the timing of biomechanical symmetry breaking determined in this study follows the early activation of Wnt signaling in concert with additional position-specific factors. We hypothesize that factors, which trigger biomechanical symmetry breaking on the cellular level, are also active in the Wnt/β-catenin feedback loop of axis formation (Fig. **6**).

**Fig. 6:**
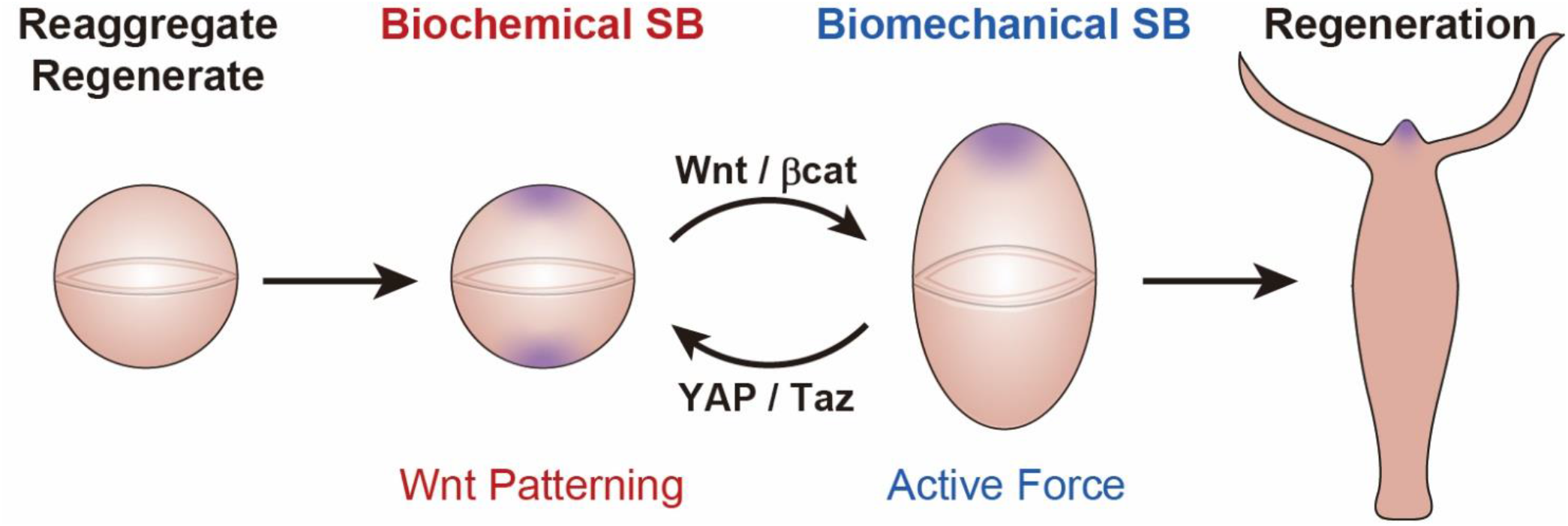
Interplays of biochemical and biomechanical symmetry breaking in *Hydra* regeneration. *Hydra* regeneration, starting from either reaggregate or regenerate, can be used as a well-defined model to unravel the interplay of biochemical and biomechanical symmetry breakings, which results in the modulation of various downstream events.

### Hippo-Yap signaling in symmetry breaking

With aid of a global transcriptome study using regenerating wildtype and β-catenin and Wnt3 overexpressing transgenic *Hydra* strains (Supporting Fig. **S3**), we identified Hippo-Yap signaling as a candidate for mechano-transduction (Fig. **5a-c**). The activation of Hippo-Yap signaling in regenerating tip tissues at *t* = 6 – 12 h (Fig. **5c**) agrees well with the activation reported during budding of *Hydra* (53) suggesting that there is a feedback mechanism between Wnt signaling and mechanical stimuli in *Hydra* head organizer formation. YAP/TAZ proteins are an integral component of the β-catenin destruction process and the depletion of YAP/TAZ has been shown to induce β-catenin/TCF transcriptional responses (47). The expression of YAP/TAZ during regeneration (*t* = 6 – 48 h) suggests that it could act as such a regulator to establish the first symmetry-breaking event in *Hydra* regeneration (Fig. **6**).

Our siYap data further indicate that Hippo-Yap signaling in *Hydra* is important for the biomechanical integrity of the tissue. siYAP treated gastric tissue pieces exhibited a doubling of the inflation-burst frequency compared to regenerating wildtype tissue (Fig. **5d,e**) and in intact animals the number of buds was increased (53). This suggests a softening of bud tissue, which otherwise has a high intercellular tension (61). Our data are related to other patterning events, e.g. the distinctive morphological phenomenon of epithelial cell extrusion where cells can be physically expelled from the tissue (68, 69). Cells that constitutively express Yap lose their planar symmetry and ultimately extrude from a previously mechanochemical homogeneous epithelial monolayer (69). In mammalian intestinal organoids, regeneration was also driven by a transient activation of YAP (58). Interestingly, this symmetry-breaking event is not uniform but depends on cell-to-cell variability of YAP activation. Only a fraction of identical cells in the initially symmetrical sphere differentiates into Paneth cells generating the stem cell niche and asymmetric crypts and villi (58). We also found that after the mosaic inactivation of YAP in Hydra polyps by single pulse of siYAP inactivation the number of buds in the budding zone further increases (data not shown).

In summary, our data on *Hydra* regeneration point to core mechanisms in symmetry breaking processes that are evolutionary conserved in metazoan evolution, from *Hydra* to mammals (53, 58).

Since our results have demonstrated the coupling of Wnt signaling and active deformation on the symmetry breaking phenomenon, incorporating positive feedback loops between these components may provide a generalized theoretical framework for many developmental and regenerative processes in nature. Also, in the context of evolution, our findings hint at the possibility that such spontaneous symmetry breaking could well be a strategy for early multicellular animals. The uniqueness of the *Hydra* system, which enables access to both molecular and mechanical processes simultaneously, sets a quantitative basis for further development of our understanding of symmetry breaking phenomena for various biological processes and forms of life.

## ACKNOWLEDGEMENTS

This work was supported by the German Science Foundation (DFG) to T.W.H. (FG1036 and SFB1324/A05 to T.W.H. and M.T. and SFB1324/B07 to S.Ö.), the JSPS (JP22H03939 to R.S. and JP19H05719 and JP20H00661 to M.T.), seed fund of Mechanobiology Institute (T.H.), and the Excellence Clusters Cell Networks (T.W.H.) and 2082/1−390761711 (M.T.). We thank Takao Ohta for stimulating discussions and Yukio Nakamura for the preparation of the actin::HyWnt3 pBSSA-AR vector. M.T. thanks Nakatani Foundation for support.

## MATERIALS AND METHODS

### Cultures and sample preparation

*Hydra magnipapillata* and b-catenin overexpressing *Hydra vulgaris* were cultured in modified *Hydra* medium (HM), containing 1mM Tris, 1mM NaHCO_3_, 0.1mM KCl, 0.1mM MgCl_2_, 1mM CaCl_2_, at 18 °C (70). The culture was synchronized. Animals were fed daily with *Artemia* and cleaned a few hours after feeding. Prior to the experiments, *Hydra* were starved for 24 h.

Reaggregates were prepared by mechanically dissociating adult *Hydra* in hypotonic dissociation medium (DM) containing 3.6 mM KCl, 6 mM CaCl_2_, 1.2 mM MgSO_4_, 6 mM sodium citrate, 6 mM pyrovic acid, 4 mM glucose, 12.5 mM TES, and 50 mg/L Rifampicin (Wako, Osaka, Japan) (7). After 2 h, the reaggregate was transferred to a glass bottom dish (MatTek Corporation, MA, US) for imaging. The medium was exchanged from DM to HM step-wisely.

Regenerates were prepared by cutting out a part of the gastric region with a scalpel under a stereoscope (SZ2-ST, Olympus, Tokyo, Japan). A piece of the tissue rolls up to form a hollow sphere in approximately 2 h. The regenerate was transferred to a glass bottom dish filled with HM.

### Imaging and image analysis

Active deformation and motion of reaggregates and regenerates were monitored using an IX81 inverted microscope (Olympus, Tokyo, Japan) every 15 s. Live imaging with high temporal resolution allowed us not only to monitor the active deformation but also to capture the moment of burst accurately. The captured images were binarized and analyzed with a self-written algorithm in MATLAB (MathWorks, MA, US).

To evaluate the directional order of actin cytoskeletons, reaggregates and regenerates were relaxed with 2 % urethane, fixed with 4 % paraformaldehyde, and stained with Alexa-fluor 488 Phalloidin (1:600, Invitrogen, CA, US) prior to the imaging using a confocal fluorescent microscope (A1R, Nikon, Tokyo, Japan). The order parameter was calculated using a self-written algorithm in MATLAB.

### Theoretical model

Biomechanical symmetry breaking is theoretically modeled by assuming *Hydra* reaggregates and regenerates as ellipsoids. The model equations are given by the minimization of a potential under a fixed volume condition, 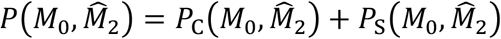, which consists of contributions of the principal curvature *P*_C_ and the surface area *P*_S_, given by

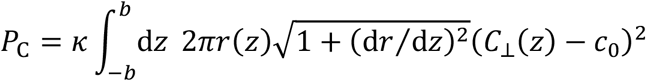

and

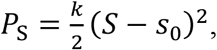

respectively. Here, *z* is the coordinate along the major axis of the ellipsoid (Fig. **1g**). The radius of the ellipsoidal section circle at the coordinate *z* is represented by *r*(*z*). The maximum principal curvatures on the ellipsoid at the coordinate *z* is denoted by *C*_⊥_(*z*). There are five parameters in this model: the constants representing bending elasticity per area and spontaneous curvature, *κ* and *c*_0_, respectively, the actual surface area *S*, the optimal area *s*_0_, and the coefficient to express the rigidity against the area change from this optimum *k* (see Supporting Information **S2** for details). We adopted the following values for these parameters: The bending rigidity is set to be 1 nJ, as reported by Naik et al. (31). The area rigidity is set to be 0.77 × 10^6^ N/m^3^, which is consistent with the value expected from the previous report (17) in the order of magnitude. See Supporting Information for how these values were extracted. The optimal surface areas are calculated using *M*_0_ = 270 and 160 μm for reaggregate and regenerate cases, which reflects the minimum values in Fig. **1d,f**, respectively. The anisotropic spontaneous curvature (*c*_0_) is altered over time as given in Fig. **1h,i**. The fixed volume is changed over time, which mimics the growth process of *Hydra* reaggregate / regenerate.

### *In situ* hybridization and transcriptomics

Whole-mount *in situ* hybridization was performed as described previously (71). RNA from biological triplicates of regeneration tip specific tissue were isolated and prepared for RNAseq as described by Petersen et al (26). RNA from Wnt3 overexpressing AEP animals (Act::Wnt3-Act::RFP) and RNA from β-catenin overexpressing animals (14) was isolated by the same protocol. The regeneration specific data are available at NCBI Hydra 2.0 Genome Project Portal (Petersen Trinity Transcripts).

## Notes

### Competing Interest Statement

The authors have declared no competing interest.

