## Supplementary Information for "Spatio-temporal Coordination of Active Deformation Forces and Wnt / Hippo-Yap Signaling in *Hydra* Regeneration"

**Supporting Information S1: Derivation of the mechanical model**

*Hydra* reaggregates and regenerates are approximated by a rotationally symmetric ellipsoid as shown in Fig. **1g**. The variables $M_0$, $M_2$ and $\widehat{M}_2$ are defined in the main text using the projected shape in two dimensions: The zero-th mode $M_0$ represents the average of the minor and major axis half-lengths $a$ and $b$ of the projected ellipse, respectively. The second mode $M_2$ represents the deviation from the circle $M_2 \equiv (a-b)/2$ , and $\widehat{M}_2 \equiv M_2/M_0$ is the degree of elliptical shape, i.e. for $\widehat{M}_2 = 0$, the 2D projection is a circle whereas for $\widehat{M}_2 = 1$, it is an infinitely thin ellipse with the length $2M_0$.
In this model, we assume that the actual *Hydra* shape is given by $\left(M_0, \widehat{M}_2\right)$ which minimizes the energy cost

$$P\left(M_0, \widehat{M}_2\right) = P_C\left(M_0, \widehat{M}_2\right) + P_S\left(M_0, \widehat{M}_2\right), \tag{1}$$

under the fixed volume condition $V = V_0$, where $V$ is the actual volume of the ellipsoid and $V_0$ is a given constant volume. Here, $P_C$ and $P_S$ are the energy costs corresponding to curvature elasticity and area elasticity of the ellipsoid surface, respectively. These two contributions are given in the following: The first term $P_C$ represents the curvature energy cost, for which an anisotropic spontaneous curvature is assumed. Specifically, we assume that the spontaneous curvature acts only in the direction of the maximum principal curvature on the ellipsoid. Then, the equation is given as

$$P_C = \kappa \int_{-b}^{b} \mathrm{d}z \; 2\pi r(z) \sqrt{1 + (\mathrm{d}r/\mathrm{d}z)^2} \left(C_\perp(z) - c_0\right)^2. \tag{2}$$

Here, $z$ is the coordinate along the major axis of the ellipsoid (Fig. **1g**), and $\kappa$ and $c_0$ are the constants representing bending elasticity per area and spontaneous curvature, respectively. Let $r(z)$ represent the radius of the ellipsoidal section circle at the coordinate $z$. The maximum principal curvatures on the ellipsoid at the coordinate $z$ is denoted by $C_\perp(z)$, which is given by

$$C_\perp(z) = a^{-1}\left[1 - \left(1 - \frac{a^2}{b^2}\right)\left(\frac{z}{b}\right)^2\right]^{-1/2}. \tag{3}$$

The circumference of the circle of the cross section at the coordinate $z$ is described as $2\pi r(z)$, and $\sqrt{1 + (\mathrm{d}r/\mathrm{d}z)^2}$ arises due to the nonlinear geometrical correction in the surface area located between the coordinates $z$ and $z$ + d$z$. Note that the other principal curvature is given by

$(a/b)^2 C_\perp(z)$, which is indeed always smaller than $C_\perp(z)$ by the definition of $a$ and $b$.

The second term $P_S$ represents the energy cost when the surface area deviates from the optimum, given by

$$P_S = \frac{k}{2}(S - s_0)^2, \tag{4}$$

where $S$ is the actual surface area, $s_0$ is the optimal area and $k$ is the coefficient to express the rigidity against the area change from this optimum.

We minimize the total potential $P(M_0, \widehat{M}_2)$ under the fixed volume condition. For this manipulation, there are a few technical notes: Firstly, by manually performing the integral in Eq. (2), the analytical expression of $P_C$ is obtained as

$$P_C = 2\pi\kappa \left\{ c_0 M_0 \left[ c_0 M_0 \left(1 - \widehat{M}_2\right)^2 - 4\left(1 + \widehat{M}_2\right)\right] + \frac{1+\widehat{M}_2}{1-\widehat{M}_2}\left[2 + c_0^2 M_0^2 \left(1 - \widehat{M}_2\right)^2\right] \frac{\arcsin\sqrt{1-\left(\frac{1-\widehat{M}_2}{1+\widehat{M}_2}\right)^2}}{\sqrt{1-\left(\frac{1-\widehat{M}_2}{1+\widehat{M}_2}\right)^2}} \right\} \tag{5}$$

Secondly, using the standard formulae for the volume and surface area of an ellipsoid, the volume $V$ and surface area $S$ are given as the functions of $M_0$ and $\widehat{M}_2$ by

$$V = \frac{4\pi}{3} a^2 b = \frac{4\pi}{3} M_0^3 \left(1 - \widehat{M}_2\right)\left(1 - \widehat{M}_2^2\right), \tag{6}$$

and

$$S = 2\pi a^2 \left(1 + \frac{b}{ae}\arcsin e\right) = 2\pi M_0^2 \left(1 - \widehat{M}_2\right)^2 \left[1 + \frac{(1+\widehat{M}_2)^2}{2(1-\widehat{M}_2)\sqrt{\widehat{M}_2}} \arcsin\left(\frac{2\sqrt{\widehat{M}_2}}{1+\widehat{M}_2}\right)\right], \tag{7}$$

respectively. Lastly, through the fixed volume condition $V(M_0, \widehat{M}_2) = V_0$ with Eq. (6), $M_0$ is implicitly given by the function of $V_0$ and $\widehat{M}_2$, and hence the potential minimum is searched with a fixed $V_0$ over $\widehat{M}_2$ from $\widehat{M}_2 = 0.0$ to $\widehat{M}_2 = 1.0$ by definition.

**Supporting Information S2: Parameters used in the model**

We used the parameter values (see **Supplementary Table S1** for list of parameter values) determined as follows: For bending rigidity, we used the value $\kappa = 1$ nJ, given by Naik et al. (1). The spontaneous curvature $c_0$ is a variable in our study. For optimal surface area, we used the value $R_0$ = 270 (reaggregate) and 160 (regenerate) μm, which are the minimum values shown in Fig. **1d,f**. The optimal volume $V_0$ is the controlled parameter, which mimics the growth of *Hydra* reaggregate / regenerate, for our purpose.

The rigidity coefficient for the area change $k$ was evaluated in the following way: Trushko et al. (2) defined the compressional rigidity $\lambda$ in terms of energy potential, given by

$$E_\lambda = \lambda \int_{L_0} \Gamma(l)^2 \, \mathrm{d}l, \tag{8}$$

where $l$ represents the arc length, and $\Gamma(l)$ represents the local extension rate in length of a cell (MDCK) layer. It is important to note that the difference of the definition of $\lambda$ from our area rigidity $k$. Trushko et al.(2) assumed a $2d$ thin elastic ring with circumferential length $L_0$, and the integration is not an areal integration but a circumferential integration. Hence, their rigidity $\lambda$ is defined over length and has the unit of [force / length]. On the other hand, our area rigidity $k$ is defined as

$$P_S = \frac{k s_0^2}{2} \cdot \frac{(s - s_0)^2}{s_0^2}, \tag{9}$$

and $k s_0^2/2$ has the unit of [force × length]. Therefore, we convert these two parameters through the relation

$$\frac{k s_0^2}{2} = \lambda L_0^2. \tag{10}$$

For this conversion, we assume $L_0$ is the typical perimeter of a section, $L_0 = 2\pi R_0$ in our case.

Since $\lambda$ is the value for a layer of MDCK cells, we substitute it with a similar tissue compressional rigidity of *Hydra* $\hat{E}$ = 10 – 150 × $10^{-3}$ N/m , which is defined $\hat{E} = Eh/2(1-\mu^2)$ with Young's modulus $E$, tissue thickness $h$ and Poisson's ratio $\mu$. For simplicity, we applied the middle value $\lambda = 80 \times 10^{-3}$ N/m with $L_0 = 2\pi R_0$ and $s_0 = 4\pi R_0^2$ with $R_0$ = 160 μm (i.e. the above-mentioned initial $R_0$

value) into Eq. (10), obtaining $k = 1.56 \times 10^6$ N/m$^3$. We used a range of $k$ considering the broad range of $\hat{E}$. In particular, the results shown in **Supplementary Information S3** used either this value of $k$ or half of it. Moreover, Fig. **1h,I** of the main text was plotted with the half value; see **Supplementary Information S3** for more details.

Furthermore, we set the force and length units, denoted by $F$ and $L$, based on the bending rigidity $\kappa$ and the reference size $R_0$. In what follows, we adimensionalize the parameters by the units $F = \kappa L^{-1} = 5.56 \times 10^{-6}$ N and $L = R_0$. The optimal surface area is $4\pi$ by the definition of the length unit assuming the shape is close to a perfect sphere right after the burst. Spontaneous curvature $c_0$ is presumed to be around 1.0 ($c_0 = 1.0 - 2.0$ are examined below) since the sphere forms spontaneously. For the regenerates $R_0 = 160$ µm, the area rigidity coefficient $k$ is adimensionalized to either 1.6 or 0.8. For the reaggregate $R_0 = 270$ µm, $k \sim 0.58$ in the adimensionalized form, thus we use $k$ which is in the value range for reaggregates and regenerates.

**Supporting Information S3: Calculation of the model**

**Supplementary Figure S2** plots the resultant relation between $M_0$ and $\widehat{M}_2$, and investigates the dependencies on the parameters $c_0$ and $\kappa$. The curves have dependency on $k$ as well as $c_0$. Firstly, we plotted $\widehat{M}_2$ against $M_0$ for various $c_0$ for $k$ = 1.6 (**Supplementary Figure S2a**) and $k$ = 0.8 (**Supplementary Figure S2b**). For both cases, we found curves that were convex upward. For increasing $c_0$, the maximum $\widehat{M}_2$ increased. The degree of decrease in $\widehat{M}_2$ with respect to increase in $M_0$ is weaker for $k$ = 0.8. Fig. **S2c** compares our theoretical curve and experimental observation (for regenerate: Fig. **1f**). The slope and magnitude of $\widehat{M}_2$ for $k$ = 0.8 (**Supplementary Figure S2b**) agree quantitatively well with experimental data. Therefore, we adopted the parameter value $k$ = 0.8 for the results shown in the main text (Fig. **1h,i**).

**Supporting Information S4: Comparative transcriptome analysis of regenerating head tissue and Wnt3 / β-catenin overexpressing polyps**

Based on a proteome/-transcriptome analysis of Hydra head regeneration (3), we established a list of all transcripts that were regulated during head regeneration and compared them with the transcriptome of *β-catenin* and *Wnt3* overexpressing animals. It included a total of 15,244 sequences that were analyzed by DESeq2. **Supplementary Figure S3** shows the heat map of all transcripts that were significantly regulated under at least one condition (padj < 0.05). By hierarchical clustering, we identified 15 clusters that were based on the similarity of their regulatory patterns. There was a sharp boundary between clusters during early regeneration (0.5 – 12 h) and late regeneration (12 – 48 h). Many transcripts encoding for cell communication and signal transduction were upregulated upon regeneration, particularly all members of the Wnt pathways (cluster 2) became activated concomitantly with the onset of regeneration. Some clusters reveal differences between *β-catenin* and *Wnt3* overexpressing animals, indicating differences in the regulation of Wnt signaling between *β-catenin* and *Wnt3* overexpressing animals.

The analysis of genes in cluster 1 revealed that genes of the Hippo-Yap pathway were upregulated during head regeneration (see also Fig. **5**). Although animals overexpressing *Wnt3* or *β-catenin* show differences in gene regulation, all members of the Hippo-Yap and Wnt pathway were strongly enriched up to 48 h after onset of regeneration. These data clearly emphasize the importance of Hippo-Yap signaling for *Hydra* regeneration.

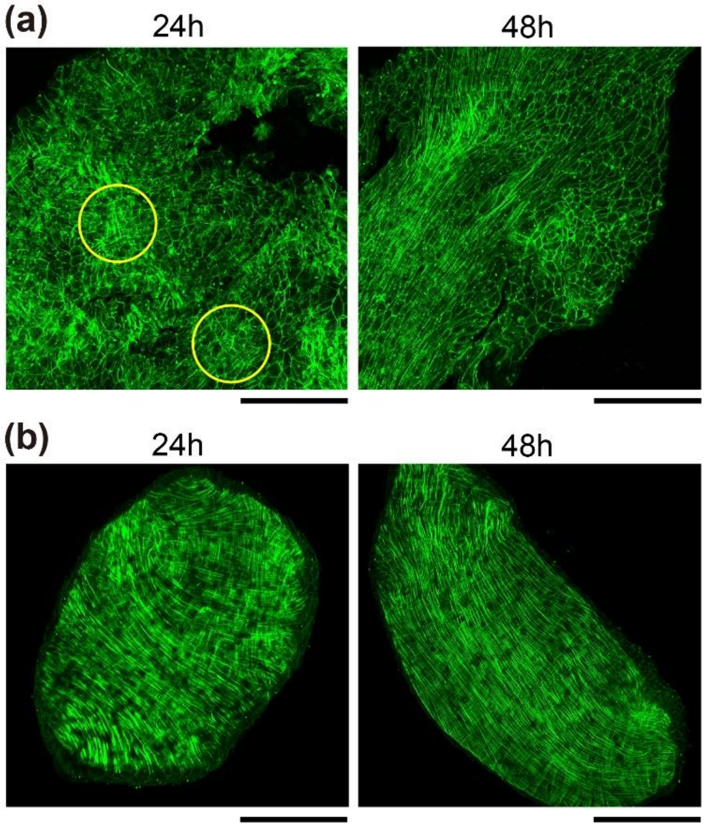


**Supplementary Figure S1: Temporal change in actin structure during regeneration.** (a) A reaggregate shows patches of mesh-like actin structures (yellow circle) at 24h. With time, the mesh-like structures spread across the whole tissue and after symmetry breaking occurs, an adult actin structure is regained. (b) Mesh-like actin structure is inherited by the adult *Hydra* and is maintained throughout the regenerating process. Scale bars: 300µm.

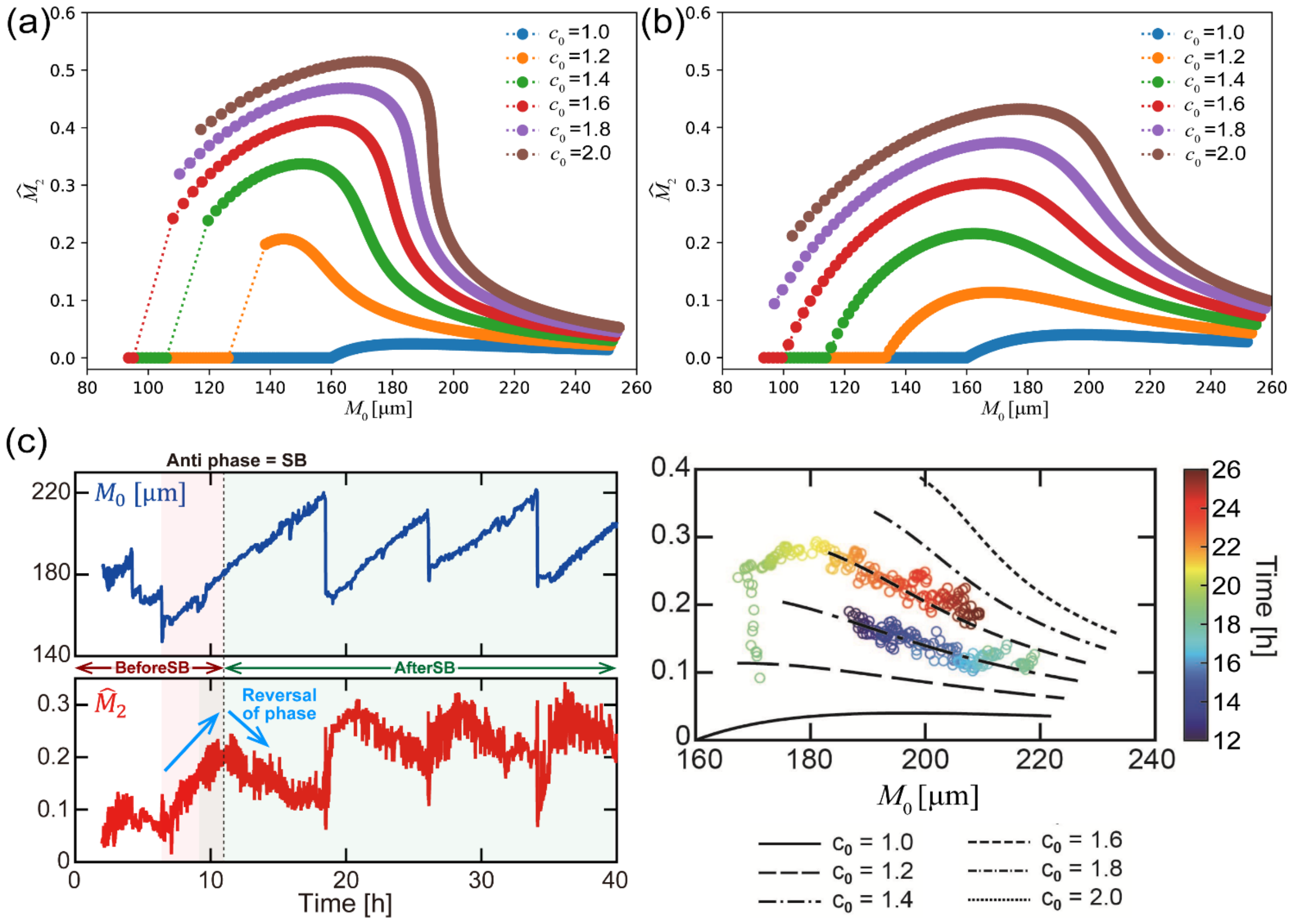


**Supplementary Figure S2: Dependence of parameters $V_0$, $c_0$ and $k$ on $M_0$ and $\widehat{M}_2$ in the mechanistic model.** (a, b) Theoretical result for (a) $k = 1.6$ and (b) $k = 0.8$. Spontaneous curvature $c_0$ ranges from 1.0, which matches the inverse radius for the sphere case, to 2.0. The colors are specified in the inset legends. The points with the same color correspond to various values of $V_0$; from left to right, $V_0$ is increased. (c) Comparison between model and experimental data for the regenerate case. Here, we used $k = 0.8$.

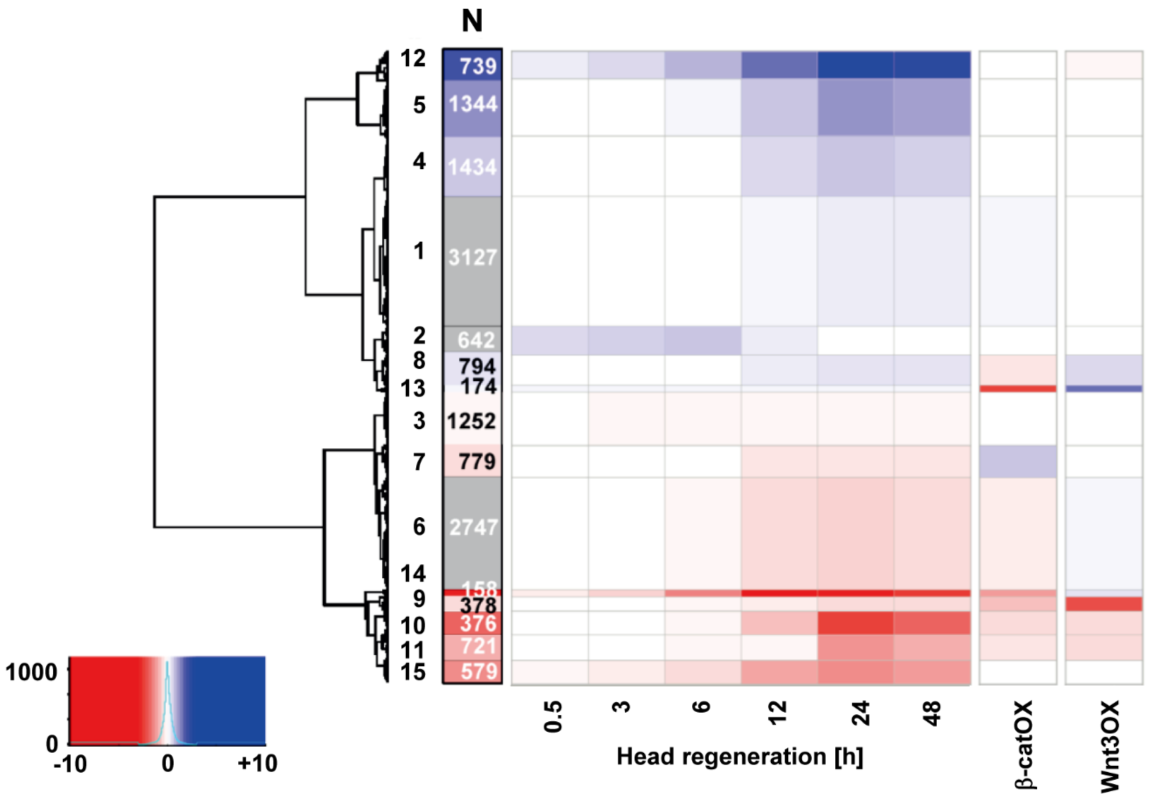


**Supplementary Figure S3: Heat map of average expression levels.** Heat map of average expression levels in regeneration tip tissue at the side of head formation and in *Wnt3* and *β-catenin* overexpressing shown by log2-fold changes. Insert indicates upregulated (blue) and downregulated genes strains (red). 12 = cell communication and Wnt signaling, 5= cell communication and transport, 4 = signal transduction including Ras/Raf, 8 = cell organization and transcription factors, 13 = cytoskeleton and DNA mismatch repair, 3 = RNA metabolism, 7 signal transduction and cell communication, 14 = regulation of gene expression and translation, 9 and 11 = proteolysis and protein metabolism, 10 biosynthesis and metabolism, 15 signal transduction and protein folding). Note the upregulation of Wnt signaling in early regenerates (cluster 12) and of other pathways including members of Hippo-Yap signaling after 12 h in later regenerates (cluster 1).

**Supplementary Table S1: List of used parameter values**

| **Parameters** | **Values** | **Sources** |
|---|---|---|
| Bending rigidity $\kappa$ | 1 nJ | (1) |
| Spontaneous curvature $c_0$ | Variable | |
| Area rigidity $k$ | $0.8 \times 10^6$ N/m$^3$ | (4) |
| Optimal surface $s_0$ (reaggregate) | $4\pi$ (270 µm)$^2 \approx 9.2 \times 10^5$µm$^2$ | Our experimental result |
| Optimal surface $s_0$ (regenerate) | $4\pi$ (160 µm)$^2 \approx 3.2 \times 10^5$µm$^2$ | Our experimental result |
